# A deep reinforcement learning approach to reconstructing quaternary structures of protein dimers through self-learning

**DOI:** 10.1101/2022.04.17.488609

**Authors:** Elham Soltanikazemi, Raj S. Roy, Farhan Quadir, Jianlin Cheng

## Abstract

Predicted interchain residue-residue contacts can be used to build the quaternary structure of protein complexes from scratch. However, only a small number of methods have been developed to reconstruct protein quaternary structures using predicted interchain contacts. Here, we present an agent-based self-learning method based on deep reinforcement learning (DRLComplex) to build protein complex structures using interchain contacts as distance constraints. We rigorously tested the DRLComplex on two standard datasets of homodimeric and heterodimeric dimers (the CASP-CAPRI homodimer dataset and Std_32 heterodimer dataset) using both true and predicted contacts. Utilizing true contacts as input, the DRLComplex achieved a high average TM-score of 0.9895 and 0.9881 and a low average interface RMSD (I_RMSD) of 0.2197 and 0.92 on the two datasets, respectively. When predicted contacts are used, the method achieves the TM-score of 0.73 and 0.76 for homodimers and heterodimers respectively. The accuracy of reconstructed quaternary structures depends on the accuracy of contact predictions. Compared with other optimization methods of reconstructing quaternary structures from interchain contacts, DRLComplex performs similarly to an advanced gradient descent method and better than a Markov Chain Monte Carlo simulation method and a simulated annealing-based method. The source code of DRLComplex is available at: https://github.com/jianlin-cheng/DRLComplex

## Introduction

In cells a protein chain folds into a 3D shape called tertiary structure. The tertiary structures of two or more protein chains often interact to form a protein complex whose conformation is called quaternary structure. Proteins also often bind with other molecules (called ligands) to perform their functions (Dhakal et al. 2022). As the function of a single protein chain or a protein complex is largely determined by its three-dimensional (3D) structure, predicting protein structure from sequence is important for studying the function of proteins. However, the determination of protein quaternary structures using experimental techniques such as x-ray crystallography is still time consuming and expensive and can be only applied to a small portion of proteins. Accurate computational prediction of protein structure from sequence has become a holy grail of computational biochemistry, drug design, biophysics, and bioinformatics over the last few decades.

Like in other prediction fields such as image processing, computer vision, and natural language processing, deep learning has driven substantial progress in protein tertiary structure prediction in the recent years (Jones and Kandathil 2018; Adhikari, Hou, and Cheng 2018; Li et al. 2019; Senior et al. 2019; Yang et al. 2020; Wu et al. 2021), which has been demonstrated in the biennial Critical Assessment of Protein Structure Prediction (CASP) (Kryshtafovych, Monastyrskyy, and Fidelis 2014; Kryshtafovych et al. 2019; Won et al. 2019). A major breakthrough was made in the CASP14 by AlphaFold2(Jumper et al. 2021), which can predict atomic high-accuracy tertiary structures for most proteins.

Despite the notable progress in tertiary structure prediction, the prediction of quaternary structure of protein complexes is still in the early stage of development (Zeng et al. 2018; J. Hou et al. 2020; Quadir, Roy, Soltanikazemi, et al. 2021; Quadir, Roy, Halfmann, et al. 2021, 2; Yan and Huang 2021; Roy et al. 2021; Xie and Xu 2021). The methods for quaternary structure prediction can be subdivided into two categories: ab-initio methods(Lyskov and Gray 2008; Pierce et al. 2014; Quadir, Roy, Soltanikazemi, et al. 2021; Evans et al. 2021; Park et al. 2021) and template-based methods(Tuncbag et al. 2012; Guerler, Govindarajoo, and Zhang 2013). The template-based approach generally searches protein structure databases for similar templates for a target and predicts the quaternary structure of the target based on the template structures. While this approach works well if suitable templates are available, it does not work for most protein complexes that do not have templates.

Different ab initio methods based on the energy optimization and the machine learning prediction of interchain contacts have been developed, which have been tested in the Critical Assessment of Protein Interaction (CAPRI) experiments (Lensink et al. 2021). For instance, RosettaDock is a popular docking tool that leverages Markov Chain Monte-Carlo (MC) based algorithms for energy minimization to dock proteins (Lyskov and Gray 2008). Zdock (Pierce et al. 2014) uses a combination of various potential scoring criteria to select the quaternary structure models generated by the Fourier transformation. Recently, a gradient descent structure optimization method - GD (Soltanikazemi et al. 2022) and a simulated annealing method - ConComplex (Quadir, Roy, Soltanikazemi, et al. 2021) have been developed to build quaternary structures from predicted interchain residue-residue contacts and tertiary structures of protein chains. Due to the recent development of more accurate deep learning methods for interchain contact prediction such as DRCon(Roy et al. 2021) and GLINTER(Xie and Xu 2021), these approaches can generate high-quality quaternary structures for some proteins. The latest significant advance in the field is the extension of the end-to-end deep learning method of predicting tertiary structures to quaternary structure prediction(Baek et al. 2021; Jumper et al. 2021). For instance, AlphaFold-multimer(Evans et al. 2021) adapts the deep learning architecture of AlphaFold2 to take as input the concatenated multiple sequence alignments of monomers in multimers and the template structures if available to predict their quaternary structures. It performs better than existing docking methods on two benchmarks. However, AlphaFold-multimer can only accurately predict the quaternary structures of a portion of protein complexes. More computational methods need to be explored and developed to advance the field.

In this work, we present a novel approach based on deep reinforcement learning (DRLComplex) to generate quaternary structures of protein dimers using predicted interchain contacts and known/predicted tertiary structures as input. The work is inspired by the huge success of the deep reinforcement learning method of AlphaGo(Silver et al. 2016) in playing GO games, defeating the world GO champion multiple times. The deep reinforcement learning also achieved similar success in playing other games such as Atari(Mnih et al. 2015) and AlphaStar(Vinyals et al. 2019). Though reinforcement learning has been commonly used in gaming and robotics, it was rarely used in bioinformatics(Wang et al. 2018; H. Hou et al. 2019; Bocicor, Czibula, and Czibula 2011) and has not been applied to protein complex structure modeling.

Like AlphaGO and Atari, our approach treats a protein complex modeling process as a game and builds the quaternary structure of a protein complex (specifically a protein dimer in this work) by adjusting the position of one protein tertiary structure (ligand) with respect to another one (receptor) in the complex to find quaternary structural conformations close to the native (true) structure, guided by the predicted interchain contacts. Compared to other optimization methods like GD, Markov Chain Monte Carlo simulation (MC) and simulated annealing in ConComplex of reconstructing quaternary structures from interchain contacts, DRLComplex achieves the state-of-the-art performance.

## Results and Discussions

We compare DRLComplex with four existing methods namely GD (gradient descent)(Soltanikazemi et al. 2022), MC (Markov Chain Monte Carlo simulation), Equidock(Ganea et al. 2022) and CNS(Brünger et al. 1998; Brunger 2007) (the simulated annealing simulation used in ConComplex)(Quadir, Roy, Soltanikazemi, et al. 2021) that constructs quaternary structures. It is worth noting that Equidock is an equivariant neural network method of directly predicting quaternary structures of dimers without using interchain contacts as input. Like DRLComplex, GD, MC, and CNS all reconstruct quaternary structures from interchain contacts. The methods are evaluated on two standard protein dimer datasets: CASP_CAPRI and Std_32. The CASP_CAPRI dataset consists of 28 homodimer(Yan and Huang 2021) targets and the Std_32 contains 32 heterodimer targets(Soltanikazemi et al. 2022). It is worth noting that only 31 out of the 32 targets of the Std_32 are used because one target (1IXRA_1IXRC) has no interchain contact between its ligand and receptor.

The methods are tested in three scenarios: the optimal scenario, the suboptimal scenario and the realistic scenario. In the optimal scenario true interchain contacts are extracted from the native quaternary structures of protein dimers and used as input to guide the assembly of the true tertiary structures of monomers in the bound state into quaternary structures. An interchain contact is a pair of residues from the two chains in a dimer in which the shortest distance between their heavy atoms is less than or equal to 6Å(Quadir, Roy, Halfmann, et al. 2021; Roy et al. 2021). In the suboptimal scenarios, the predicted interchain contacts together with the true tertiary structure of monomers in the bound state are used as input. We used two interchain contact predictors: DRCon (6Å version) to predict interchain contacts for the CASP_CAPRI homodimer dataset and Glinter(Xie and Xu 2021) for the Std_32 heterodimer dataset. In the most realistic scenario both the predicted interchain contacts and the unbound tertiary structures of monomers predicted by AlphaFold2(Jumper et al. 2021) are used as input.

The performance of these methods is evaluated by comparing their reconstructed structures with the native/true structures of the dimers using TMalign(Zhang and Skolnick 2005) and DockQ(Basu and Wallner 2016). The evaluation metrics include TM-Score, root-mean-square deviation (RMSD) of the entire structure with respect to the native structure, the fraction of the native interchain contacts in the reconstructed structures (f_nat), interface RMSD (I_RMSD), and Ligand RMSD (L_RMSD).

### Performance on Std_32 Heterodimer Dataset

The performance of all the methods on the Std_32 heterodimer dataset is reported in **Table 1, Table 2** and **Table 3** for the optimal, suboptimal and the realistic scenarios, respectively. The detailed target-wise results of DRLComplex are shown in **Table S1, Table S2** and **Table S3**, respectively. In the optimal scenario (**Table 1**) the DRLComplex method outperforms all the methods in all the evaluation metrics except F_nat. Particularly, the difference in terms of RMSD is more pronounced. For instance, the RMSD is 0.88 Å, 2.9 Å, 3.1 Å and 10.04 Å for the DRLComplex, GD, MC and CNS, respectively. In the suboptimal scenario (**Table 2**), DRLComplex’s performance is overall better than the other method. For instance the TMscore of the DRLComplex is 1.33%, 4.1%, 11.7% and 24.5% better than the GD, MC, CNS and the Equidock respectively.

**Table 1.** Average mean of RMSD, TM-score, f_nat, I_RMSD, and L_RMSD results of DRLComplex, GD, MC, and CNS on the Std_32 dataset with true interchain contacts and true tertiary structure of monomers as inputs. Bold numbers denote the best results.

| Methods | TM-score | RMSD | F_nat (%) | I_RMSD | L_RMSD |
| --- | --- | --- | --- | --- | --- |
| DRLComplex | **0.98** | **0.88** | 90.03 | **0.92** | **2.15** |
| GD | 0.95 | 2.9 | **92.43** | 1.99 | 7.16 |
| MC | 0.94 | 3.1 | 92.24 | 2.2 | 7.18 |
| CNS | 0.82 | 10.04 | 69.13 | 3.71 | 14.99 |

**Table 2.** Average mean RMSD, TM-score, f_nat, I_RMSD, and L_RMSD of the models reconstructed by the DRLComplex, GD, MC, CNS and Equidock on Std_32 with predicted interchain contacts and true tertiary structure of monomers as inputs. Bold numbers denote the best results.

|  | TM-score | RMSD | F_nat (%) | I_RMSD | L_RMSD |
| --- | --- | --- | --- | --- | --- |
| DRLComplex | **0.76** | 13.93 | **19.64** | **13.68** | **34.16** |
| GD | 0.75 | **13.92** | 16.68 | 13.72 | 34.17 |
| MC | 0.73 | 14.14 | 13.88 | 13.82 | 35.36 |
| CNS | 0.68 | 17.09 | 17.26 | 15.81 | 43.12 |
| Equidock | 0.61 | 18.53 | 4.95 | 14.98 | 36.11 |

**Table 3.** Average mean RMSD, TM-score, f_nat, I_RMSD, and L_RMSD of the models reconstructed by the DRLComplex, GD, MC, CNS and Equidock methods on Std_32 with predicted interchain contacts and predicted tertiary structures of monomers as inputs. Bold numbers denote the best results.

| Methods | TM-score | RMSD | F_nat (%) | I_RMSD | L_RMSD |
| --- | --- | --- | --- | --- | --- |
| DRLComplex | **0.74** | **14.53** | **11.80** | 13.86 | 36.18 |
| GD | 0.73 | 14.54 | 11.71 | **13.85** | **36.17** |
| MC | 0.7 | 15.06 | 11.25 | 14.60 | 36.66 |
| CNS | 0.66 | 19.02 | 3.78 | 14.74 | 37.91 |
| Equidock | 0.59 | 18.63 | 3.55 | 14.42 | 35.97 |

Similarly, in the realistic scenario (**Table 3**), the DRLComplex’s performance is overall better as it has the best performance in TM-score, RMSD, F_nat. As seen in Table 6, the TM-score of the DRLComplex is 1.3%, 5.7%, 12.2% and 25.4% better than the GD, MC, CNS and the Equidock respectively. **Figure 1** illustrates the reconstruct quaternary structures for a heterodimer in comparison with the true structure in the three scenarios.

**Figure 1.**
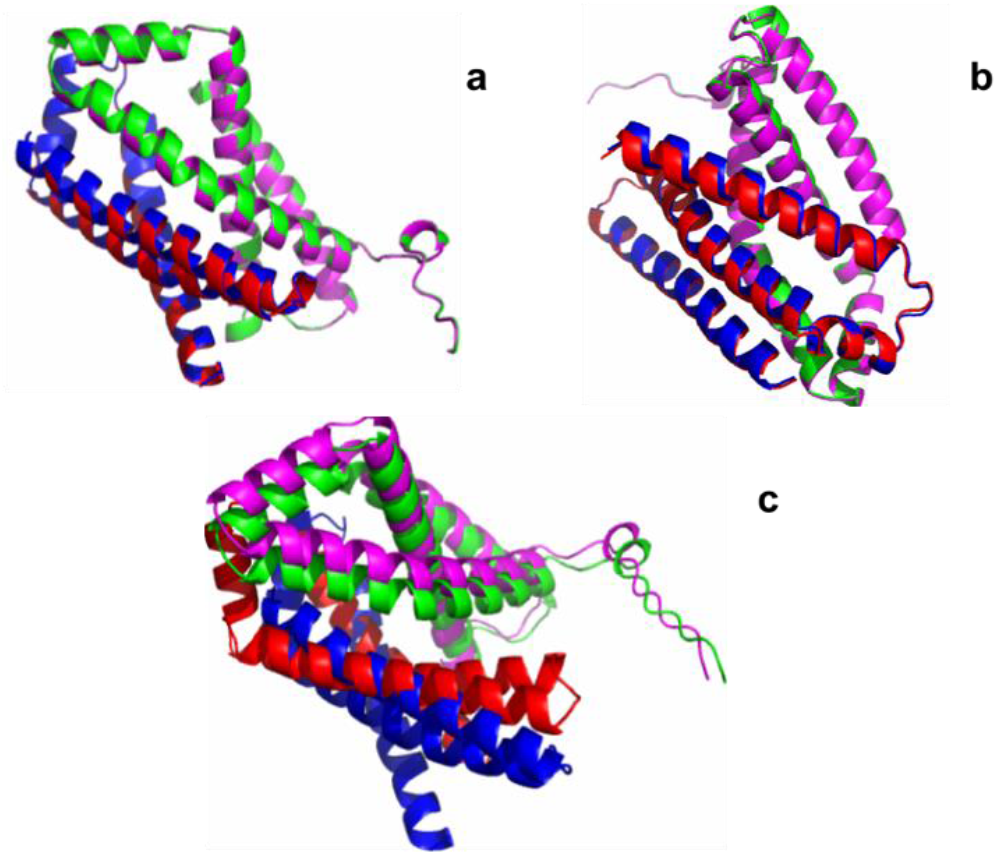
The superposition of the native structure of a heterodimer (PDB code: 2WDQ) and the models reconstructed by DRLComplex (i.e., green and red: true structure of the two chains in the dimer; blue and magenta: predicted structure of the two chains): (a) using true tertiary structures and true interchain contacts as inputs, (b) using true tertiary structures and predicted interchain contacts as input, (c) using predicted tertiary structures and predicted interchain contacts as input. TM-score, RMSD, f_nat, I_RMSD, L_RMSD of the model predicted by DRLComplex for (a) are 0.97, 0.94Å, 97.1%, 0.95 Å, and 1.85Å, respectively, for (b) are 0.99, 0.65Å, 92.1%, 0.58 Å, and 1.12Å, respectively, and for (c) are 0.75, 6.03Å, 1%, 6.07Å, and 10.6Å, respectively.

### Performance on CASP_CAPRI Homodimer Dataset

The average performance of the methods on the CASP_CAPRI homodimer dataset for the optimal scenario, suboptimal scenario and the realistic scenario is reported in **Table 4, Table 5** and **Table 6** respectively. The detailed per-target results of the DRLComplex on the three different settings are shown in **Table S4, Table S5** and **Table S6**. In the optimal scenario (**Table 4**), DRLComplex performs equally well or better than all other predictors in terms of all the evaluation metrics, followed by GD. For 10 out of the 28 targets, DRLComplex reconstructs quaternary structures with a perfect f_nat score of 1. The average fraction of true interchain contacts recalled in the reconstructed structures (f_nat) is 99.05%, indicating highly accurate quaternary structures are reconstructed by the DRLComplex for the dimers when the optimal input is provided. Equidock was not included in this scenario as it does not use any interchain contacts as input, but rather directly predicts the translation and rotation of the ligand with respect to the receptor.

**Table 4.** Mean of RMSD, TM-score, f_nat, I_RMSD, and L_RMSD of the DRLComplex, GD, MC and CNS on 28 homodimers on the CASP-CAPRI dataset using true interchain contacts and true tertiary structures as inputs. Bold numbers denote the best results.

| Methods | TM-score | RMSD | F_nat (%) | I_RMSD | L_RMSD |
| --- | --- | --- | --- | --- | --- |
| DRLComplex | **0.9895** | **0.3753** | **99.05** | **0.2197** | **0.8235** |
| GD | **0.9895** | **0.3753** | 99.03 | 0.3468 | **0.8235** |
| MC | 0.9631 | 1.2089 | 78.91 | 1.1611 | 2.8897 |
| CNS | 0.9234 | 2.003 | 73.45 | 3.9234 | 4.6841 |

**Table 5.** Mean of RMSD, TM-score, f_nat, I_RMSD, and L_RMSD of the DRLComplex, GD, MC, CNS, and Equidock on 28 homodimers in CASP-CAPRI dataset using predicted interchain contacts and true tertiary structures of monomers as inputs. Bold numbers denote the best results.

| Methods | TM-score | RMSD | F_nat (%) | I_RMSD | L_RMSD |
| --- | --- | --- | --- | --- | --- |
| DRLComplex | 0.73 | 11.88 | 33.21 | 10.43 | 26.52 |
| GD | **0.74** | **11.09** | **35.51** | **9.66** | **25.21** |
| MC | 0.72 | 12.04 | 32.65 | 10.47 | 26.98 |
| CNS | 0.62 | 14.55 | 28.32 | 14.37 | 36.25 |
| Equidock | 0.56 | 18.57 | 6.51 | 14.5 | 35.24 |

**Table 6.** Mean of RMSD, TM-score, f_nat, I_RMSD, and L_RMSD of the DRLComplex, GD, MC, CNS and Equidock on 28 homodimers in CASP-CAPRI dataset using predicted interchain contacts and predicted tertiary structures of monomers as inputs. Bold numbers denote the best results.

| Methods | TM-score | RMSD | F_nat (%) | I_RMSD | L_RMSD |
| --- | --- | --- | --- | --- | --- |
| DRLComplex | 0.64 | 12.18 | 27.1 | 10.73 | 26.54 |
| GD | **0.69** | **12.15** | **28.05** | **10.69** | **26.50** |
| MC | 0.63 | 12.78 | 26.81 | 11.89 | 28.92 |
| CNS | 0.62 | 14.55 | 13.58 | 12.62 | 31.69 |
| Equidock | 0.50 | 26.59 | 22.27 | 18.39 | 44.66 |

In the sub-optimal scenarios (**Table 5**), GD, DRLComplex and MC’s performance is close, while they outperform CNS and Equidock by a large margin. For instance, the average TM-score of the DRLComplex method is 17.7% and 30.4% higher than the CNS and the Equidock respectively.

Furthermore, in the realistic scenario (**Table 6**), GD performs best and DRLComplex the second. DRLComplex performs better than the other methods. For instance, the TM-score of the complexes constructed by the DRLComplex is 1.59%, 3.22% and 28% better than the MC, CNS and Equidock respectively. It is noticed that the TM-score of DRLComplex drops from 0.989 to 0.73 because the predicted contacts contain fewer true contacts and some false contacts.

**Figures 2, 3**, and **4** illustrates how the quality (TM-score and f_nat) of the reconstructed complex structures changes with respect to the precision, recall and F1-score of the predicted contacts. It is shown that the quality of the reconstructed structures largely increases with the increase of the accuracy of predicted interchain contacts.

**Figure 2.**
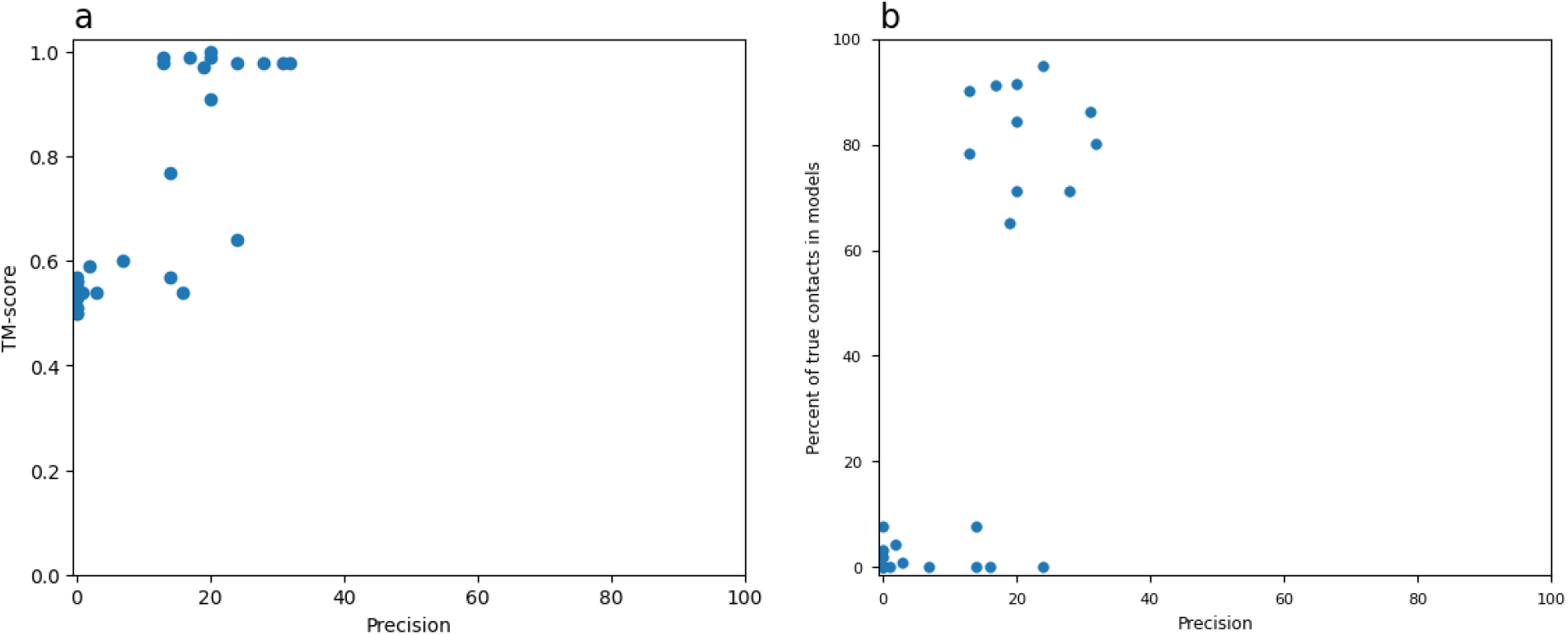
TM-score and F_nat of the models generated by DRLComplex VS the precision of the predicted interchain contacts on the CASP-CAPRI homodimer dataset. The true tertiary structures of monomers are used as input.

**Figure 3.**
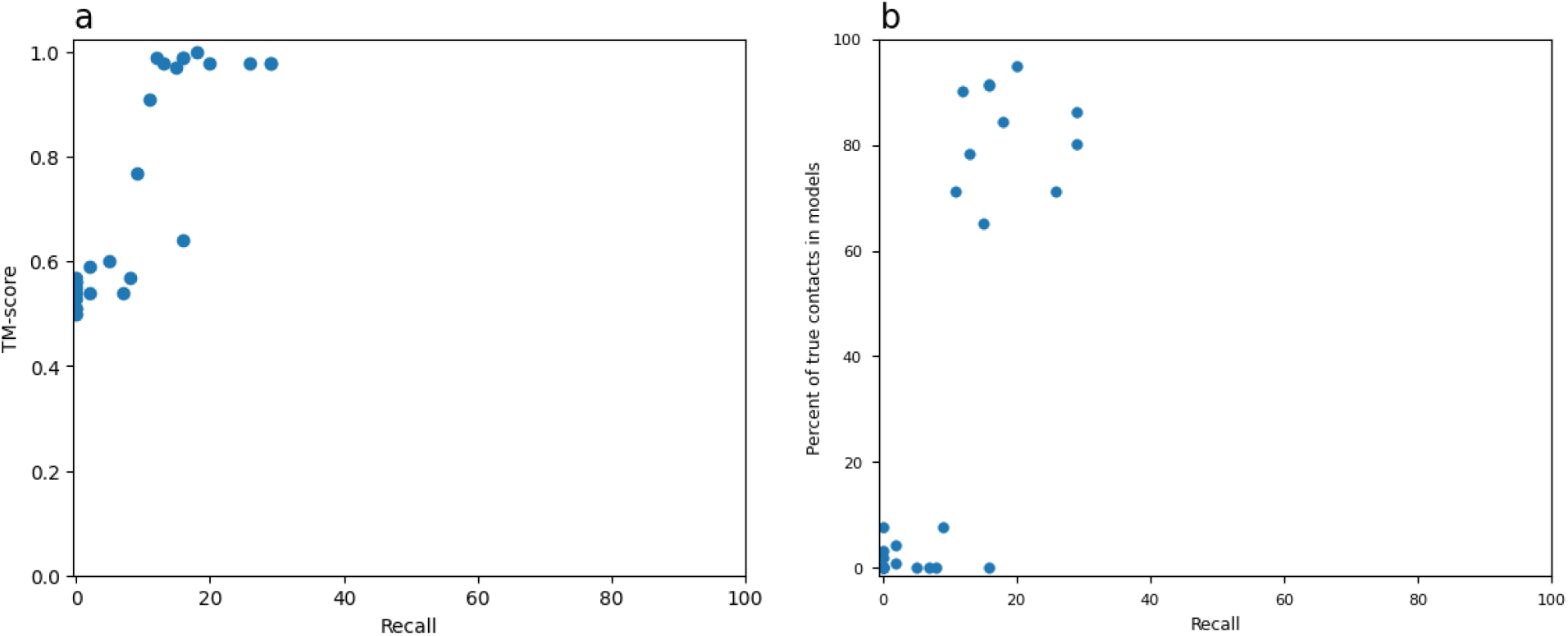
TM-score and F_nat of the models generated by DRLComplex VS the recall of the predicted interchain contacts on the CASP-CAPRI dataset. The true tertiary structures of monomers are used as input.

**Figure 4.**
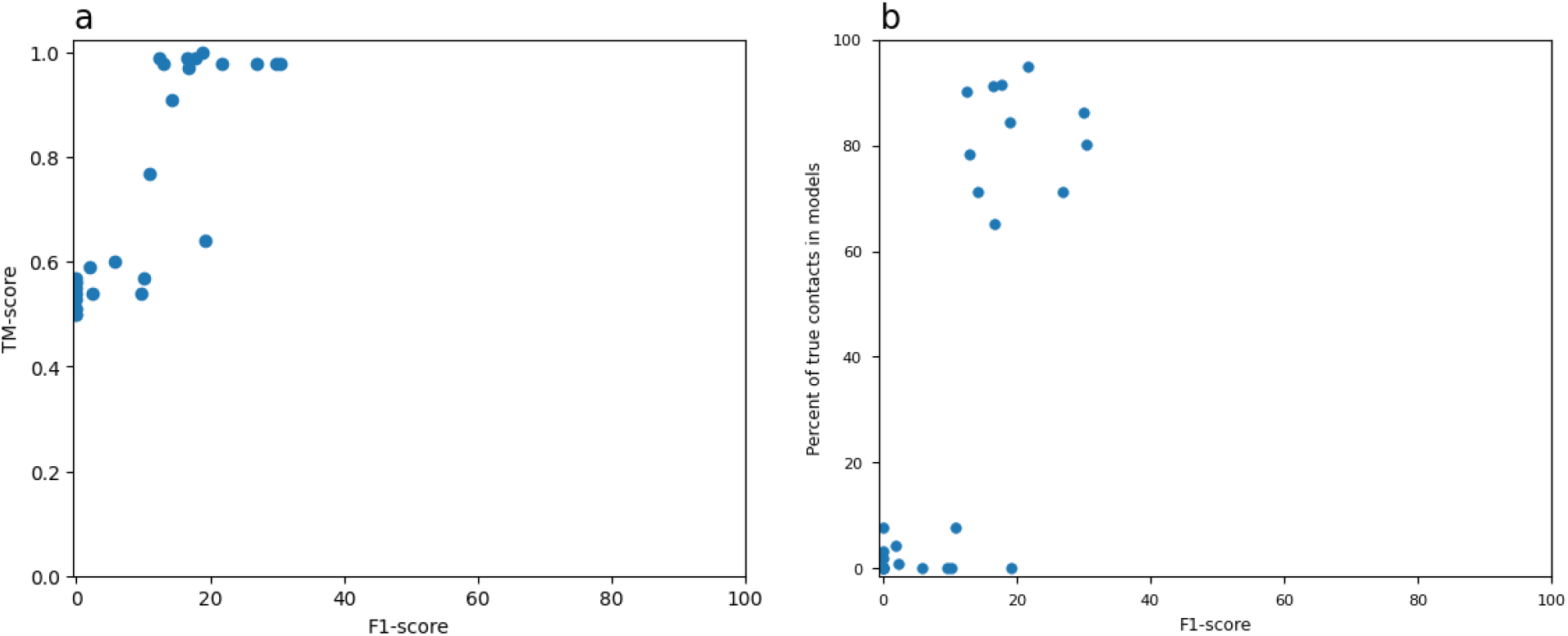
TM-score and F_nat of the models generated by DRLComplex VS the F1-score of the predicted interchain contacts on the CASP-CAPRI dataset. The true tertiary structures of monomers are used as input.

## Materials and Methods

DRLComplex uses deep reinforcement learning techniques to automatically construct the quaternary structure of protein dimers by adjusting the position of one protein chain (ligand) with respect to another one held fixed (receptor) to produce structural conformations similar to the native (true) structure. Specifically, an artificial intelligence (AI) *agent* is trained to choose *actions* according to both *intermediate reward* and *long-term reward* to modify the *state* (structure) of a protein dimer in a modeling *environment*. The state (*S*) of a protein dimer is represented by two equivalent forms: the 3D coordinates of atoms and the interchain distance map. The former is suitable for generating new structures in the environment by applying actions to it and the latter is suitable for a deep learning model to predict the values of actions. The interchain distance map is represented by a 2D matrix (*M*) with a size of *L*_1_ × *L*_2_, where *L*_1_ is the length of the ligand and *L*_2_ is the length of the receptor. The cells of the matrix hold the distances between *C*_*β*_ atoms of the residues from the ligand and the receptor in a dimer.

We use six actions including three translations with step size of 1Å and three rotations with the step size of 1° along the x-, y-, and z-axis (forward and backward) to adjust the position of one chain (the ligand) against the other one (the receptor). A deep convolutional neural network (Q) approximating the Q-function of the reinforcement learning is used to predict the Q-value (i.e., an estimate of the immediate and future rewards) of the six actions (**Figure 5**) on an input state *S*. The input to the deep neural network is the interchain distance map (*M*) representing *S*. This deep network is called by the agent to predict the values of the actions, which are used to choose the next action to change the state of a dimer.

**Figure 5.**
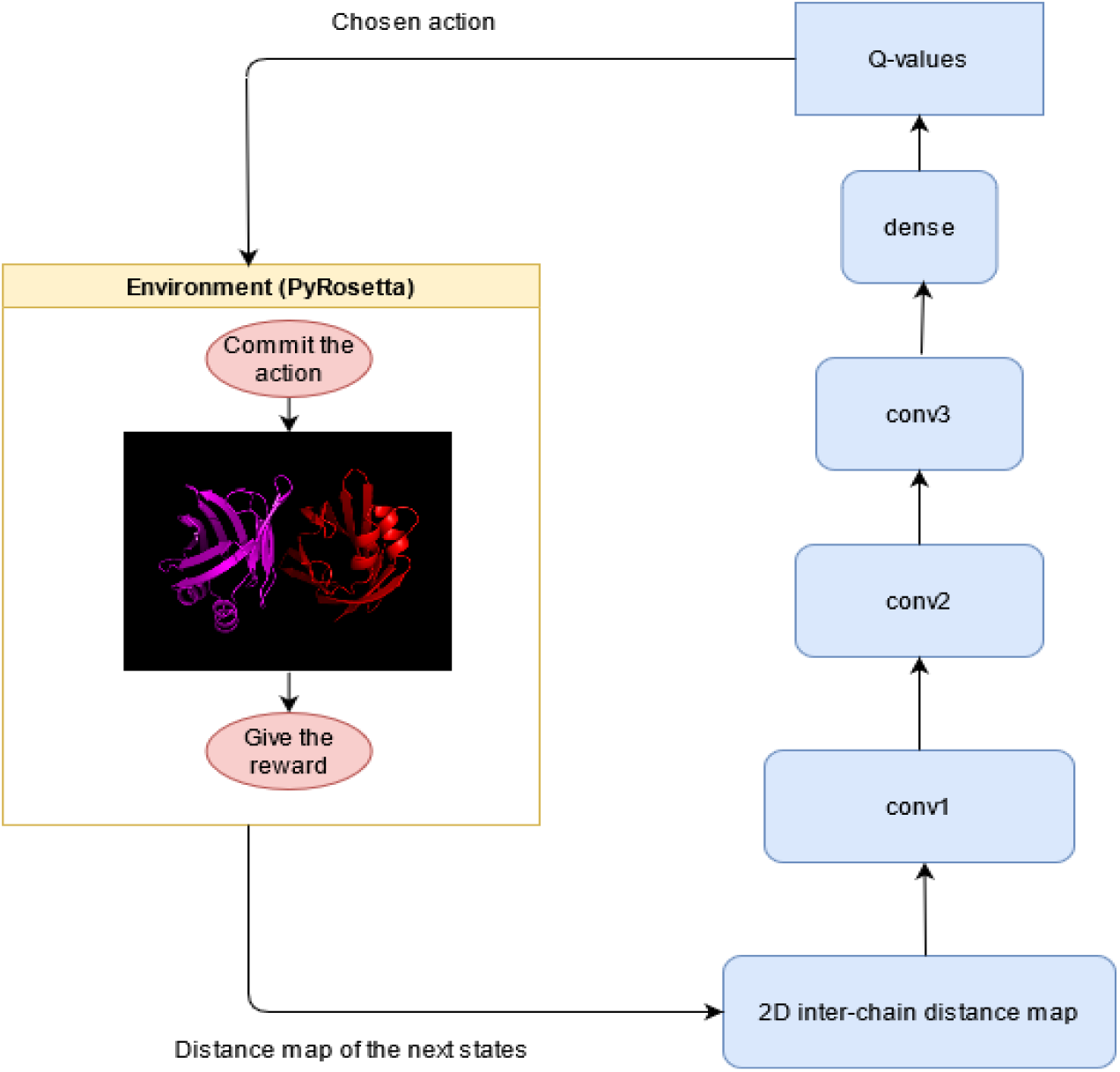
The deep network to predict the values of six actions given the interchain distance map (S) of the state of a dimer as input. The first convolutional layer has 16 filters of size 3 × 3 with stride 2. The second layer contains 32 filters with stride 2, and the filter size is 5 × 5. The third convolutional layer has 64 3 × 3 filters with stride 2. The rectified linear unit (ReLU) function is used by all the three convolutional layers. The output of the third convolutional layer is then fed into a fully connected layer to generate hidden features, which are used by the fully connected output layer with the linear activation function to predict Q-values of the six actions.

The deep convolutional neural network is trained by a sequence of examples (state (*S*), action (*A*), reward (*R*), next State (*S’*)), which are automatically obtained by the agent continuously interacting with the modeling environment implemented by PyRosetta to choose actions to adjust the position of the ligand according to a *ϵ* − *greedy* policy. Given a state *S* at each step (episode), the agent selects either an action having the highest value predicted by the deep network with a probability of 1 − *ϵ* or a random action with *ϵ* probability. The action (A) is then applied to the 3D structure of the current state S by the environment (E) to generate a new 3D structure of the next state S’. The immediate reward (R) of the action is calculated as the difference between the quality of S’ and S with respect to the target state (S*), which can be either the true structure of the dimer or the predicted interchain contact map provided to the agent at the beginning. The process is repeated to generate many examples. We use a technique called experience replay to train the deep network, which stores the agent’s experience at each step (S, A, R, S’) in a buffer. Training with experience replay is illustrated in **Figure 6**.

**Figure 6.**
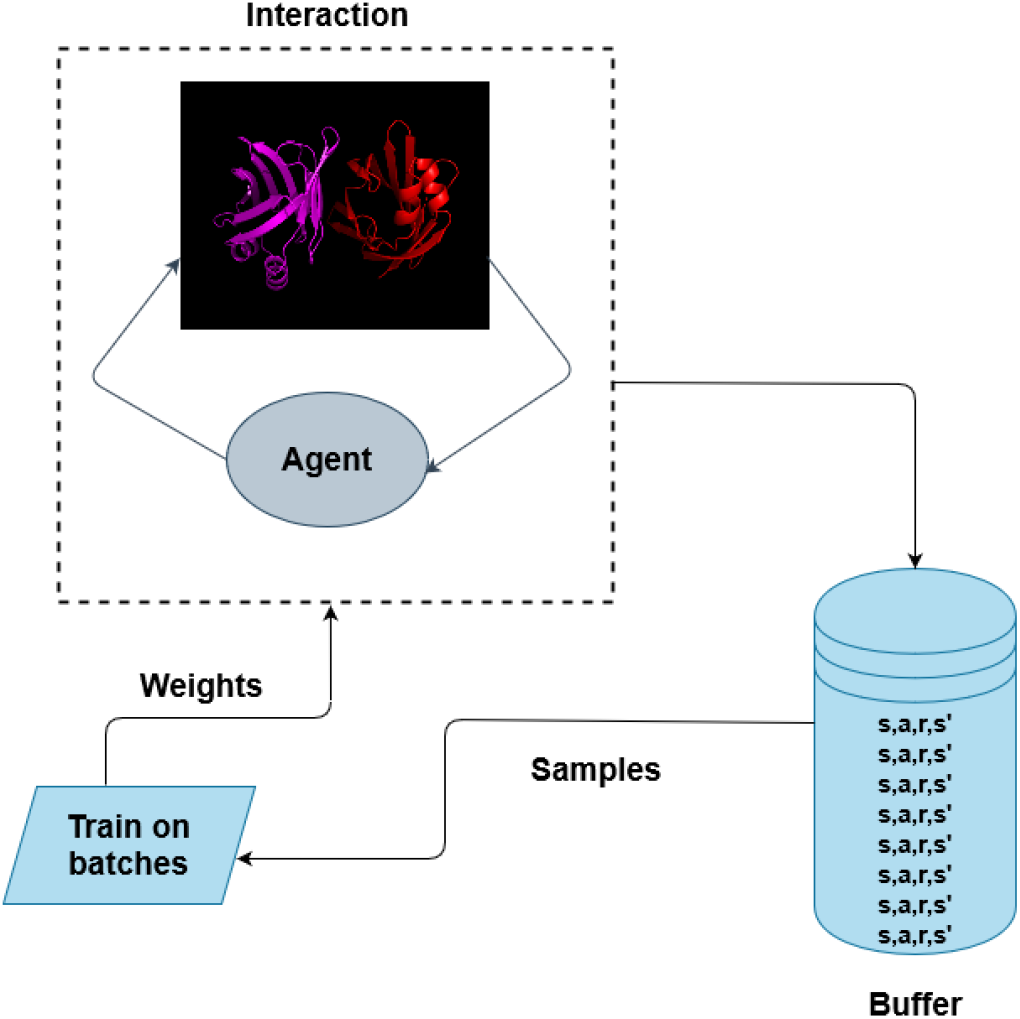
Training of the deep network for a protein dimer with the experience replay. The agent interacts with the environment to generate a sequence of experiences/examples (<s, a, r, s’> vectors). The experiences are stored in a buffer. If the buffer is full, some earliest experiences are deleted to release space for new experiences. K transition experiences are sampled from the buffer to train the deep network (i.e., updating the Q-value function of reinforcement learning). The process is repeated until the system converges (i.e., a final state is reached).

We use the immediate reward and the output of a so-called target network (Q_target_) to generate the reference Q-value (Q_reference_(S, a)) for an action *a* and a state *S* as the labels to train the deep network (Q) of the agent. The target network uses a copy of the weights of the deep neural network of the agent at every k steps to predict the Q-value of each action *a*′ (*Q*_*target*_(*s*′, *a*′)) for the next state *S*’ (**Figure 7**), which is used to estimate the future value of each action. The reference Q-value for S and a (i.e., *Q*_*reference*_(*s, a*)) is the immediate reward r(S, a) plus the predicted highest future value multiplying a discount factor *γ* (i.e., *Q*_*reference*_(*s, a*) = *r*(*s, a*) + *γ*. *max*_*a*′_*Q*_*target*_(*s*′, *a*′)). The maximum future value is the maximum Q-value predicted by the target network for the next state S’ and any possible action a’. The loss (L) used to train Q is the average squared error 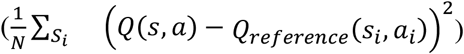, where N is the number of states. We use the stochastic gradient descent with minibatch size of 64 to train the network. During the experiment, *ϵ* of the *ϵ*-policy is decreased from 1 to 0.1 for the first 100000 iterations via the exponential decay (i.e., decay factor = 0.99) and is fixed at 0.1 thereafter.

**Figure 7.**
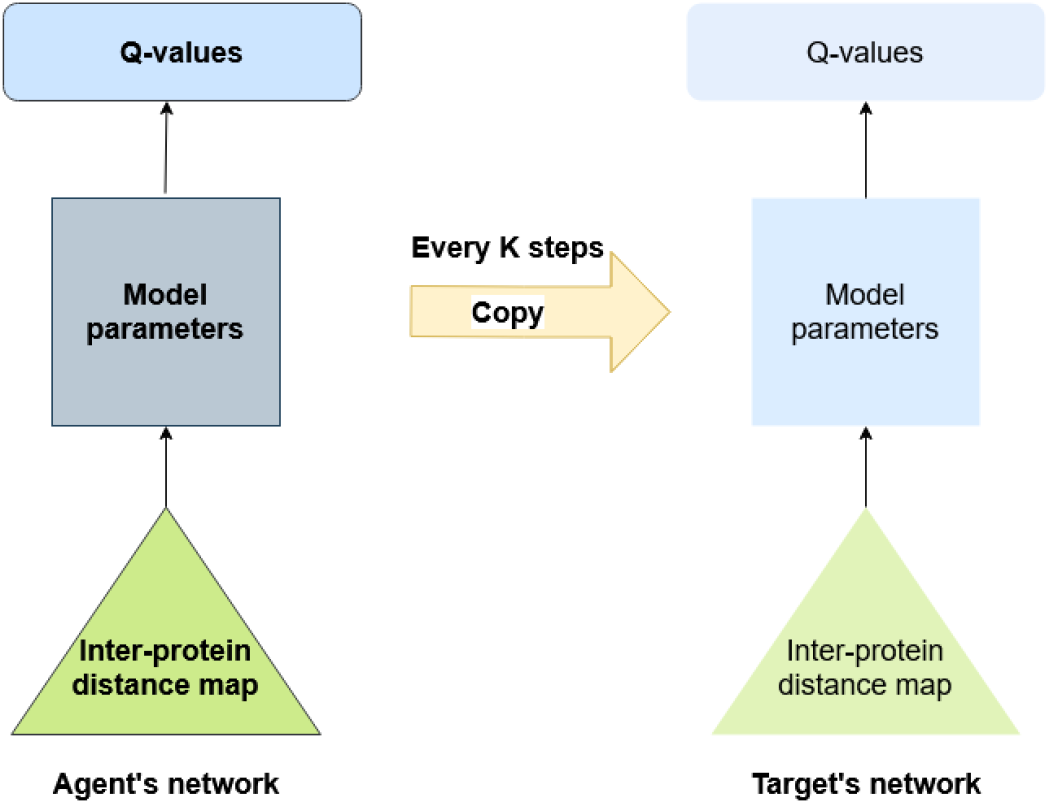
The agent’s deep network (Q) and the target network (Q_reference_) to generate reference Q-values of actions to train Q.

We test two different strategies to evaluate the quality of a state (S) to calculate the immediate reward: (1) the root mean square distance (RMSD) between the 3D structure of S and the true structure (S*), and (2) the contact energy – the agreement between the distance map of S and the interchain contact map provided as the input at the beginning. It is noted that only the latter is used in the final experiments since the true structure is typically unknown in real applications. For the first strategy, the RMSD of the current structure of S and the next state (S’) is calculated as RMSD_S_ and RMSD_S’_ and the immediate reward r(S, a) is equal to RMSD_S_ - RMSD_S’_. If the quality of the next state generated by action a is better (i.e., RMSD_S’_ < RMSD_S_), the agent receives a positive reward; otherwise, it gets a negative reward (penalty).

For the second strategy, the immediate reward is the contact energy of S minus the contact energy of S’. The contact energy function measuring the satisfaction of a contact between any two residues is defined as follows:

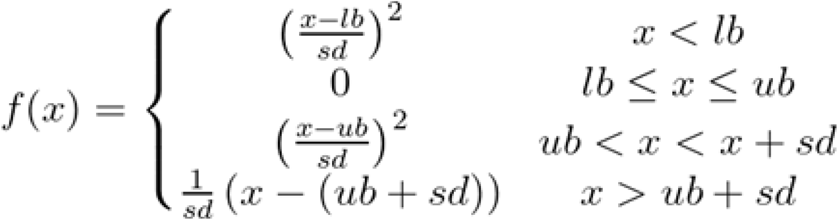

Here, *lb* and *ub* represent the lower bound and upper bound of the distance (*x*) between two residues that are predicted to be in contact. Two residues are considered in contact if the distance between their closest heavy atoms is less than or equal to 6 Å. The lower bound (*lb*) is empirically set to 0 and the upper bound (*ub*) to 6 Å. *sd* is the standard deviation, which is set to 0.1. Based on this cost function, if the distance between two residues predicted to be in contact is <= 6 Å, i.e., the contact restraint is satisfied, and the contact energy is 0. Otherwise, it is a positive value. The complete contact energy function for the structural model of a state (S or S’) is the sum of the energies for all predicted interchain contacts used in modeling (called contact energy). The immediate reward is the contact energy of (S) minus that of S’. The goal is to find actions to increase the satisfaction of contacts in the structure (i.e., to reduce the contact energy).

**Figure 8** shows how the accumulated award and the RMSD of the reconstructed structure for a protein dimer (PDB code: 1A2D) changes across episodes in the self-learning process by using the contact energy to calculate the reward. The accumulated reward converges in the last 200 episodes. The self-learning (or game) finishes by generating a structure with an RMSD of 0.94 Å. The experiment demonstrates that a high-quality structure can be generated by using deep reinforcement learning to maximize the rewards based on the predicted interchain contacts. The deep network is trained for 100K steps, using a batch size of 32. The target network is also updated every 500 steps. It is worth noting that the deep network is trained specifically for each target through self-learning. Therefore, the inference time for each target involves the training time as well.

**Figure 8.**
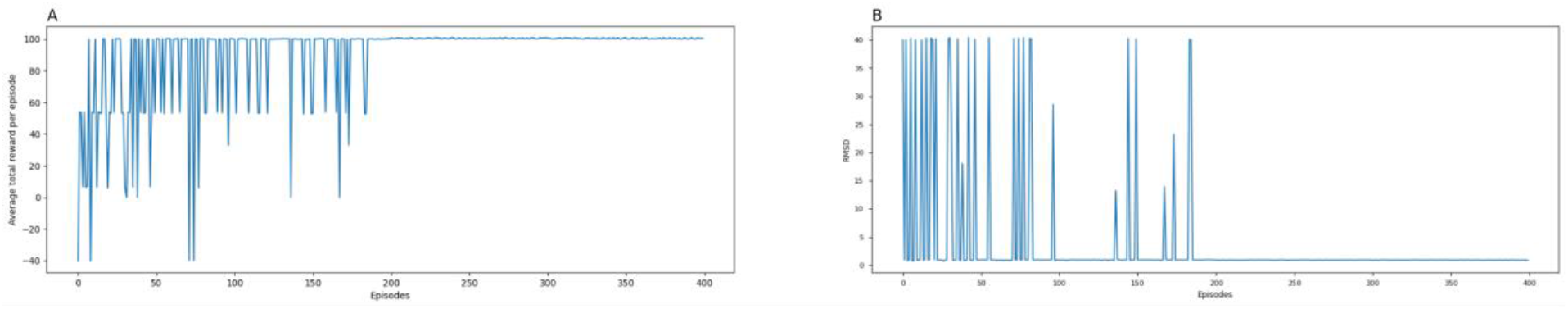
The total rewards and the RMSD of the reconstructed structures at each episode for a protein dimer (PDB code: 1A2D) using the contact energy to calculate the reward. (A) The average total reward at each episode averaged over several self-learning runs (games). (B) The RMSD of the predicted structure at the end of each episode.

## Conclusion

We develop a novel deep reinforcement learning method (DRLComplex) to reconstruct the quaternary structures from interchain contacts. It is the first such method to use the reinforcement self-learning to adjust the position of one protein (ligand) with respect to the other one (receptor) considering both short-term and long-term rewards to search for the near-native quaternary structure. Our experiments demonstrate that the deep reinforcement learning method can generate high-quality structural models for nearly all the dimers when true interchain contacts are provided. It also can construct the structural models of protein complexes from predicted contacts with reasonable quality if the input is sufficiently accurate. Testing it in different circumstances shows that it can achieve the state-of-the-art performance of reconstructing the structures of protein dimers form interchain contacts. In the future, we plan to pretrain DRLComplex on a large set of representative protein dimers so that it can directly predict the actions of assembling the two units of any new dimer into near native structures without using the on-the-fly self-learning in order to speed up the prediction process.

