## Supplementary Materials for "A deep reinforcement learning approach to reconstructing quaternary structures of protein dimers through self-learning"

**Table S1.** Detailed results (RMSD, TM-score, F_nat, I_RMSD and L_RMSD) of DRLComplex on Std_32 with true interchain contacts and true tertiary structures as inputs. The average TM-score is 0.987 with a min of 0.97 and a max of 0.99.

| **Target** | **RMSD** | **TM-score** | **F_nat (%)** | **I_RMSD** | **L_RMSD** |
| --- | --- | --- | --- | --- | --- |
| 3RRLA_3RRLB | 0.93 | 0.98 | 98.2 | 0.902 | 2.441 |
| 2NU9A_2NU9B | 0.95 | 0.99 | 96.1 | 0.995 | 1.678 |
| 1EP3A_1EP3B | 0.84 | 0.99 | 99.3 | 1.044 | 1.541 |
| 2Y69B_2Y69C | 0.74 | 0.99 | 84.1 | 0.756 | 2.28 |
| 3RPFA_3RPFC | 0.93 | 0.97 | 50 | 0.864 | 2.475 |
| 1TYGB_1TYGA | 0.81 | 0.99 | 99.2 | 0.927 | 1.986 |
| 3MMLA_3MMLB | 0.96 | 0.99 | 94.8 | 0.932 | 2.063 |
| 2VPZA_2VPZB | 0.91 | 0.99 | 93.8 | 1.172 | 2.305 |
| 2Y69A_2Y69C | 0.86 | 0.99 | 82.6 | 0.916 | 2.123 |
| 1I1QA_1I1QB | 0.87 | 0.99 | 98.8 | 0.906 | 2.069 |
| 2Y69A_2Y69B | 0.75 | 0.99 | 86.3 | 0.842 | 1.96 |
| 1EFPA_1EFPB | 0.97 | 0.99 | 97.5 | 1.04 | 3.152 |
| 1W85A_1W85B | 0.77 | 0.99 | 99.3 | 0.76 | 2.263 |
| 1ZUNA_1ZUNB | 0.93 | 0.99 | 98.11 | 0.954 | 1.761 |
| 3PNLA_3PNLB | 0.87 | 0.99 | 99.4 | 0.782 | 2.077 |
| 3OAAH_3OAAG | 0.9 | 0.99 | 93.8 | 1.008 | 1.851 |
| 3G5OA_3G5OB | 0.86 | 0.97 | 90.2 | 0.839 | 1.685 |
| 2WDQC_2WDQD | 0.94 | 0.97 | 97.1 | 0.95 | 1.853 |
| 1BXRA_1BXRB | 0.96 | 0.99 | 97.6 | 1.14 | 3.119 |
| 1RM6A_1RM6B | 0.79 | 0.99 | 96.5 | 0.604 | 1.892 |
| 1QOPA_1QOPB | 0.78 | 0.99 | 97.2 | 0.761 | 1.625 |
| 1B70A_1B70B | 0.96 | 0.99 | 92.4 | 1.123 | 2.165 |
| 3A0RA_3A0RB | 0.8 | 0.99 | 98.1 | 0.699 | 2.017 |
| 2ONKA_2ONKC | 0.9 | 0.99 | 96.4 | 0.535 | 1.932 |
| 2D1PB_2D1PC | 0.92 | 0.97 | 83.7 | 0.935 | 2.325 |
| 4HR7A_4HR7B | 0.93 | 0.99 | 82.6 | 1.03 | 2.472 |
| 1RM6A_1RM6C | 0.95 | 0.99 | 74.5 | 1.323 | 2.434 |
| 3IP4B_3IP4C | 0.93 | 0.99 | 79.7 | 0.946 | 2.628 |
| 3IP4A_3IP4C | 0.97 | 0.99 | 74.3 | 1.158 | 2.77 |
| 1RM6B_1RM6C | 0.81 | 0.99 | 69.3 | 0.9 | 1.696 |
| Mean | 0.88 | 0.987 | 90.03 | 0.92 | 2.15 |

**Table S2**. The RMSD, TM-Score, f_nat, I_RMSD, and L_RMSD of DRLComplex for 31 hetero dimers in Std_32 using predicted contacts as input. The true tertiary structures of the monomers are used in this experiment. A target which does not have any interchain contacts is discarded.

| **Target** | **TM-score** | **RMSD** | **F_nat(%)** | **I_RMSD** | **L_RMSD** |
| --- | --- | --- | --- | --- | --- |
| 1EFPA_1EFPB | 0.87 | 3.19 | 15 | 3.24 | 7.82 |
| 1EP3A_1EP3B | 0.67 | 8.08 | 11 | 6.45 | 17.9 |
| 1I1QA_1I1QB | 0.76 | 19.29 | 5 | 21.2 | 55.52 |
| 1QOPA_1QOPB | 0.62 | 20.38 | 5 | 23.72 | 51.28 |
| 1W85A_1W85B | 1 | 0.73 | 80 | 0.48 | 2.02 |
| 1ZUNA_1ZUNB | 0.65 | 15.48 | 8 | 16.54 | 30.22 |
| 2D1PB_2D1PC | 0.49 | 14.89 | 4 | 14.17 | 34.35 |
| 2NU9A_2NU9B | 0.89 | 2.63 | 48 | 2.23 | 5.17 |
| 2ONKA_2ONKC | 0.52 | 27.47 | 2 | 25.26 | 65.44 |
| 2VPZA_2VPZB | 0.84 | 34.39 | 2 | 32.21 | 92.48 |
| 2WDQC_2WDQD | 1 | 0.65 | 92 | 0.58 | 1.12 |
| 2Y69A_2Y69B | 0.68 | 26.1 | 8 | 27.41 | 65.18 |
| 2Y69A_2Y69C | 0.61 | 25.66 | 6 | 25.21 | 59.85 |
| 2Y69B_2Y69C | 0.53 | 20.94 | 10 | 16.56 | 65.58 |
| 3A0RA_3A0RB | 0.82 | 17.56 | 7 | 20.38 | 43.67 |
| 3G5OA_3G5OB | 0.66 | 8.85 | 4 | 9.41 | 13.83 |
| 3IP4A_3IP4B | 0.5 | 25.71 | 8 | 20.64 | 57.94 |
| 3IP4A_3IP4C | 0.87 | 12.22 | 7 | 14.91 | 31.5 |
| 3IP4B_3IP4C | 1 | 0.67 | 89 | 0.44 | 2.16 |
| 3MMLA_3MMLB | 0.9 | 2.64 | 27 | 2.36 | 5.62 |
| 3OAAH_3OAAG | 0.93 | 3.79 | 35 | 4.22 | 7.93 |
| 3PNLA_3PNLB | 0.7 | 6.95 | 6 | 6.29 | 17.11 |
| 3RPFA_3RPFC | 0.73 | 17.47 | 7 | 9.07 | 53.93 |
| 3RRLA_3RRLB | 0.62 | 12.91 | 5 | 12.63 | 25.17 |
| 4HR7A_4HR7B | 0.95 | 8.35 | 12 | 9.6 | 23.34 |
| 1B70A_1B70B | 0.82 | 16.77 | 2 | 20.58 | 34.72 |
| 1BXRA_1BXRB | 0.81 | 20.79 | 6 | 20.82 | 49.85 |
| 1RM6A_1RM6B | 0.74 | 26.13 | 6 | 25.87 | 59.09 |
| 1RM6A_1RM6C | 0.74 | 17.4 | 9 | 18.68 | 48.44 |
| 1RM6B_1RM6C | 0.65 | 13.13 | 4 | 12.86 | 29.88 |
| 1TYGB_1TYGA | 0.91 | 0.53 | 79 | 0.34 | 0.87 |
| Mean | 0.7574 | 13.9274 | 19.6451 | 13.689 | 34.1606 |

**Table S3.** Detailed results (RMSD, TM-score, F_nat, I_RMSD, L_RMSD, TM-score of ligand, and TM-score of receptor of DRLComplex on Std_32 with predicted interchain contacts and predicted tertiary structures as inputs. The average TM-score is 0.74 with a min of 0.5 and a max of 1.

| **Target** | **RMSD** | **TM-score** | **F_nat** | **I_RMSD** | **L_RMSD** | **TM-score of Ligand** | **TM-score of Receptor** |
| --- | --- | --- | --- | --- | --- | --- | --- |
| 1EFPA_1EFPB | 2.83 | 0.89 | 25 | 3.21 | 6.08 | 0.98 | 0.96 |
| 1EP3A_1EP3B | 5.07 | 0.86 | 16 | 4.48 | 11.12 | 0.97 | 0.98 |
| 1I1QA_1I1QB | 20.79 | 0.67 | 5 | 21.52 | 62.06 | 0.97 | 0.97 |
| 1QOPA_1QOPB | 20.22 | 0.64 | 7 | 23.35 | 50.21 | 0.97 | 0.99 |
| 1W85A_1W85B | 4.64 | 0.82 | 11 | 3.33 | 5.88 | 0.99 | 0.99 |
| 1ZUNA_1ZUNB | 16.22 | 0.67 | 3 | 17.45 | 27.81 | 0.93 | 0.95 |
| 2D1PB_2D1PC | 10.61 | 0.61 | 0 | 10.77 | 28.47 | 0.99 | 0.99 |
| 2NU9A_2NU9B | 1.92 | 1 | 52 | 1.83 | 3.41 | 0.99 | 0.97 |
| 2VPZA_2VPZB | 32.16 | 0.81 | 1 | 30.17 | 89.68 | 0.98 | 0.94 |
| 2WDQC_2WDQD | 6.03 | 0.75 | 1 | 6.07 | 10.6 | 0.96 | 0.98 |
| 2Y69A_2Y69B | 24.2 | 0.77 | 9 | 23.51 | 58.93 | 0.99 | 0.98 |
| 2Y69A_2Y69C | 19.97 | 0.79 | 1 | 21.49 | 40.44 | 0.99 | 0.99 |
| 2Y69B_2Y69C | 18.91 | 0.56 | 3 | 7.11 | 58 | 0.98 | 0.99 |
| 3A0RA_3A0RB | 21.38 | 0.6 | 0 | 20.33 | 53.29 | 0.77 | 0.90 |
| 3G5OA_3G5OB | 14.11 | 0.54 | 3 | 14.14 | 26.31 | 0.90 | 0.96 |
| 3IP4A_3IP4B | 20.31 | 0.51 | 4 | 14.58 | 81.33 | 0.99 | 0.88 |
| 3IP4A_3IP4C | 11.57 | 0.82 | 1 | 14.83 | 29.61 | 0.99 | 0.92 |
| 3IP4B_3IP4C | 5.01 | 0.95 | 34 | 2.18 | 6.99 | 0.88 | 0.92 |
| 3MMLA_3MMLB | 3.83 | 0.81 | 40 | 3.2 | 9.24 | 0.99 | 0.97 |
| 3OAAH_3OAAG | 14.53 | 0.81 | 26 | 17.55 | 26.62 | 0.59 | 0.97 |
| 3PNLA_3PNLB | 7.07 | 0.69 | 1 | 6.59 | 18.67 | 0.98 | 0.99 |
| 3RPFA_3RPFC | 16.36 | 0.64 | 10 | 7.73 | 54.69 | 0.97 | 0.96 |
| 3RRLA_3RRLB | 7.66 | 0.73 | 9 | 8.03 | 12.29 | 0.99 | 0.93 |
| 4HR7A_4HR7B | 19.39 | 0.73 | 6 | 20.02 | 57.84 | 0.90 | 0.96 |
| 1B70A_1B70B | 17.59 | 0.83 | 2 | 21.32 | 36.1 | 0.97 | 0.95 |
| 1BXRA_1BXRB | 20.93 | 0.79 | 2 | 20.99 | 50.06 | 0.99 | 0.99 |
| 1RM6A_1RM6B | 26.29 | 0.6 | 4 | 25.94 | 59.11 | 0.99 | 0.99 |
| 1RM6A_1RM6C | 17.74 | 0.85 | 0 | 18.84 | 49.08 | 0.99 | 0.98 |
| 1RM6B_1RM6C | 13.9 | 0.63 | 5 | 12.95 | 29.93 | 0.99 | 0.98 |
| 1TYGB_1TYGA | 0.4 | 0.97 | 81 | 0.41 | 0.82 | 0.90 | 0.96 |
| 2ONKA_2ONKC | 28.65 | 0.5 | 4 | 25.68 | 66.83 | 0.97 | 0.93 |
| Mean | 14.525 | 0.7367 | 11.806 | 13.858 | 36.177 | 0.950 | 0.962 |

**Table S4**. The RMSD, TM-score, f_nat, I_RMSD and L_RMSD of DRLComplex for individual targets in the CASP-CAPRI dataset using true contacts and true tertiary structure as inputs. RMSDs range from 0.12 to 3.57, with a mean of 0.375 and median of 0.235. TM-Score has values ranging from 0.768 to 0.999, with an average of 0.989. For the f_nat metric, values range from 0.96 to 1 with an average of 0.99. I_RMSD has a minimum value of 0.125, a maximum value of 2.979, an average value of 0.326 and a median value of 0.241 and lastly, the L_RMSD has values ranging from 0.261 to 7.106 with an average of 0.8 and median score of 0.533.

| **Target** | **RMSD** | **TM-score** | **F_nat** (%) | **I_RMSD** | **L_RMSD** |
| --- | --- | --- | --- | --- | --- |
| T0759 | 0.22 | 0.9980 | 100.0 | 0.224 | 0.527 |
| T0764 | 0.18 | 0.9995 | 100.0 | 0.193 | 0.391 |
| T0770 | 0.26 | 0.9992 | 98.4 | 0.262 | 0.562 |
| T0776 | 0.19 | 0.9992 | 100.0 | 0.200 | 0.475 |
| T0780 | 0.15 | 0.9995 | 97.8 | 0.157 | 0.324 |
| T0792 | 0.56 | 0.9851 | 97.7 | 0.434 | 1.393 |
| T0801 | 0.22 | 0.9993 | 99.3 | 0.256 | 0.535 |
| T0805 | 0.16 | 0.9994 | 97.4 | 0.167 | 0.338 |
| T0811 | 0.23 | 0.9991 | 98.5 | 0.230 | 0.516 |
| T0813 | 0.12 | 0.9998 | 98.6 | 0.125 | 0.261 |
| T0815 | 3.57 | 0.7680 | 100.0 | 2.979 | 7.106 |
| T0819 | 0.20 | 0.9994 | 99.2 | 0.209 | 0.424 |
| T0825 | 0.37 | 0.9967 | 100.0 | 0.326 | 0.922 |
| T0843 | 0.24 | 0.9993 | 99.4 | 0.252 | 0.532 |
| T0847 | 0.21 | 0.9989 | 100.0 | 0.218 | 0.468 |
| T0849 | 0.25 | 0.9988 | 99.2 | 0.264 | 0.537 |
| T0851 | 0.22 | 0.9995 | 99.0 | 0.222 | 0.450 |
| T0852 | 0.26 | 0.9991 | 98.6 | 0.279 | 0.636 |
| T0893 | 0.38 | 0.9852 | 99.0 | 0.278 | 0.890 |
| T0965 | 0.38 | 0.9980 | 100.0 | 0.350 | 0.804 |
| T0966 | 0.24 | 0.9994 | 100.0 | 0.272 | 0.633 |
| T0976 | 0.23 | 0.9991 | 100.0 | 0.209 | 0.492 |
| T0984 | 0.29 | 0.9993 | 99.1 | 0.227 | 0.663 |
| T0999D1 | 0.38 | 0.9983 | 98.1 | 0.370 | 0.863 |
| T0999D4 | 0.28 | 0.9986 | 100.0 | 0.265 | 0.749 |
| T1003 | 0.18 | 0.9996 | 98.8 | 0.189 | 0.393 |
| T1006 | 0.34 | 0.9941 | 96.8 | 0.347 | 0.742 |
| T1032 | 0.20 | 0.9990 | 98.2 | 0.208 | 0.433 |
| Mean | 0.3753 | 0.9895 | 99.05 | 0.2197 | 0.8235 |

**Table S5**. Detailed results (RMSD, TM-score, f_nat, I_RMSD, L_RMSD) of DRLComplex for CAPS-CAPRI dataset using predicted interchain contacts and true tertiary structure. The TM-score values range from 0.50 to 0.99, with an average value of 0.73.

| **Target** | **RMSD** | **TM-score** | **F_nat** (%) | **I_RMSD** | **L_RMSD** |
| --- | --- | --- | --- | --- | --- |
| T0976 | 18.69 | 0.54 | 0 | 17.8 | 36 |
| T0776 | 4.65 | 0.77 | 7.6 | 5.706 | 13.31 |
| T0813 | 0.98 | 0.99 | 90.1 | 0.925 | 2.366 |
| T0852 | 29.34 | 0.57 | 0 | 21.71 | 48.98 |
| T0966 | 27.26 | 0.53 | 0 | 12.55 | 62.44 |
| T1003 | 0.89 | 0.99 | 91.3 | 0.833 | 1.719 |
| T0819 | 0.62 | 1 | 84.3 | 0.631 | 1.211 |
| T0965 | 16.28 | 0.59 | 4.3 | 13.86 | 28.57 |
| T0792 | 14.57 | 0.5 | 0 | 14.93 | 40.23 |
| T0851 | 1.3 | 0.98 | 80.2 | 0.858 | 1.819 |
| T0815 | 10.52 | 0.51 | 0 | 11.9 | 32.47 |
| T0770 | 24.99 | 0.54 | 0 | 23.16 | 50.39 |
| T1032 | 18.13 | 0.54 | 0.9 | 17.07 | 29.24 |
| T0999D1 | 14.42 | 0.64 | 0 | 12.66 | 27.34 |
| T0805 | 1.07 | 0.98 | 78.4 | 1.08 | 2.326 |
| T0780 | 22.7 | 0.55 | 0 | 21.52 | 55.84 |
| T1006 | 16.05 | 0.5 | 0 | 19.34 | 50.87 |
| T0843 | 0.92 | 0.99 | 91.4 | 0.942 | 1.466 |
| T0893 | 0.85 | 0.98 | 94.8 | 0.92 | 1.94 |
| T0811 | 1.08 | 0.98 | 86.3 | 0.901 | 3.675 |
| T0984 | 39.94 | 0.56 | 1.8 | 31.43 | 72.91 |
| T0849 | 1.07 | 0.98 | 71.2 | 1.086 | 3.437 |
| T0764 | 13.7 | 0.56 | 3.2 | 12.97 | 33.29 |
| T0759 | 14.09 | 0.51 | 0 | 12.38 | 38.39 |
| T0999D4 | 2.48 | 0.91 | 71.1 | 1.661 | 10.3 |
| T0825 | 17.39 | 0.57 | 7.8 | 14.91 | 32.18 |
| T0847 | 17.08 | 0.6 | 0 | 16.69 | 55.77 |
| T0801 | 1.67 | 0.97 | 65.24 | 1.72 | 3.98 |
| Mean | 11.8832 | 0.73 | 33.2121 | 10.4336 | 26.5163 |

**Table S6**. Detailed results (RMSD, TM-score, f_nat, I_RMSD, L_RMSD and TM-score of monomer) of DRLComplex for CAPS-CAPRI dataset using predicted interchain contacts and predicted tertiary structures. The TM-score values range from 0.36 to 0.97, with an average value of 0.64. The average TM-score of the monomers predicted by Alphafold2 is 0.95.

| **Target** | **TM-score** | **RMSD** | **F_nat (%)** | **I_RMSD** | **L_RMSD** | **TM-score of the monomer** |
| --- | --- | --- | --- | --- | --- | --- |
| T1003 | 0.92 | 0.58 | 78 | 0.51 | 0.8 | 0.9964 |
| T0984 | 0.48 | 33.57 | 2 | 27.17 | 57.5 | 0.9805 |
| T0792 | 0.49 | 12.7 | 7 | 12.16 | 30.47 | 0.9585 |
| T0805 | 0.94 | 1.83 | 75 | 1.85 | 2.39 | 0.9675 |
| T0851 | 0.93 | 2.03 | 73 | 1.68 | 3.01 | 0.9687 |
| T0999D1 | 0.51 | 16.52 | 2 | 13.24 | 34.22 | 0.9708 |
| T0815 | 0.49 | 14.7 | 1 | 11.29 | 47.88 | 0.9789 |
| T0759 | 0.4 | 15.62 | 7 | 12.42 | 35.6 | 0.7475 |
| T0893 | 0.36 | 20.09 | 32 | 15.39 | 54.24 | 0.7178 |
| T0825 | 0.49 | 18.23 | 0 | 15.64 | 30.46 | 0.9655 |
| T0819 | 0.88 | 1.94 | 47 | 2.28 | 2.22 | 0.9705 |
| T1006 | 0.47 | 11.39 | 0 | 11.93 | 33.96 | 0.9859 |
| T0966 | 0.48 | 32.4 | 9 | 30.28 | 76.42 | 0.9529 |
| T0770 | 0.53 | 11.7 | 8 | 13.61 | 28.94 | 0.9777 |
| T0843 | 0.95 | 1.04 | 63 | 0.9 | 1.72 | 0.988 |
| T0999D4 | 0.59 | 7.03 | 0 | 8.09 | 17.99 | 0.9767 |
| T0780 | 0.46 | 22.17 | 4 | 21.1 | 56.72 | 0.9599 |
| T0976 | 0.52 | 18.16 | 3 | 16.99 | 34.17 | 0.9777 |
| T0811 | 0.96 | 0.88 | 77 | 0.87 | 2.24 | 0.993 |
| T0852 | 0.45 | 33.26 | 6 | 21.91 | 52.15 | 0.9334 |
| T1032 | 0.6 | 6.25 | 45 | 5.6 | 8.49 | 0.7 |
| T0776 | 0.75 | 4.8 | 26 | 3.57 | 18.46 | 0.9765 |
| T0801 | 0.88 | 2.12 | 57 | 2.3 | 4.11 | 0.9648 |
| T0813 | 0.97 | 1.14 | 69 | 1.34 | 1.2 | 0.9818 |
| T0764 | 0.55 | 15.14 | 1 | 16.74 | 27.8 | 0.9882 |
| T0849 | 0.88 | 1.38 | 64 | 1.14 | 1.79 | 0.9727 |
| T0965 | 0.57 | 15.64 | 1 | 13.24 | 27.57 | 0.9851 |
| T0847 | 0.52 | 18.81 | 2 | 17.41 | 50.72 | 0.9901 |
| Mean | 0.64 | 12.18 | 27.1 | 10.74 | 26.54 | 0.95 |

[**Supplementary Video**](https://drive.google.com/file/d/19CaJAIfDhibt_t_iLr_WnfKAJ6Vxwufx/view?usp=sharing)

A video showing DRLComplex reconstructing the quaternary structure of a dimer (PDB code: 1A2D) from true interchain contacts.

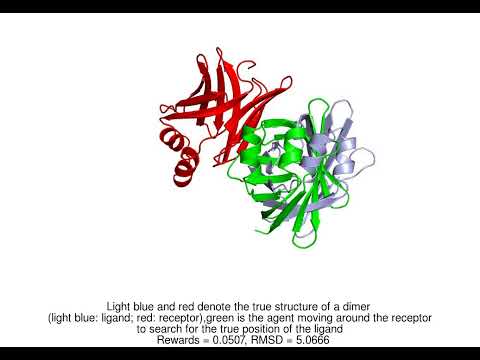
<https://www.youtube.com/embed/MXja2zSck7Q?feature=oembed>
